# Divergent realized ecological niches of sister species of the Antarctic amphipod genus *Charcotia*

**DOI:** 10.1101/2024.08.14.607880

**Authors:** Dorien Aerts, Aaron Kolder, Gilles Lepoint, Loïc N. Michel, Isa Schön

## Abstract

Climate change and resource exploitation in the Southern Ocean are important Anthropogenic pressure on Antarctic food webs. Understanding the eco-functional roles of Antarctic communities is essential for ecosystem management and conservation. Amphipods are among the most dominant and ecologically important benthic taxa in the Southern Ocean. The amphipod genus *Charcotia* is part of the scavenger guild playing a dominant role in nutrient recycling. To study the trophic habits of two sister species, *C. amundseni* and *C. obesa*, stable isotope ratios of carbon and nitrogen were investigated along geographical and bathymetrical gradients. *Charcotia obesa* belongs to the fourth and *C. amundseni* to the fifth trophic level, based on significant differences in δ^15^N. Benthic primary producers dominate the diet in both species as derived from their δ^13^C values. *Charcotia obesa*, the species with the narrowest depth range, did not show a depth-related pattern in isotopic ratios. An increasing geographic gradient of both δ^15^N and δ^13^C values was observed for *C. obesa*, ranging from the northern most tip of the Western Antarctic Peninsula to the southwestern most part of the Bellingshausen Sea. This might be linked to nutrient rich glacial meltwater in the latter part of the Southern Ocean. Nitrogen stable isotope ratios of *C. amundseni* showed a significant difference between Crown Bay and the other locations; this might be explained by the close location of the Filchner Area to nutrient rich upwelling in the Weddell Sea Gyre. Our study provides evidence for niche differentiation between two closely related amphipod species. Incorporation of additional samples from other locations and depth ranges in combination with isotope analysis and DNA-based prey identification might clarify the trophic position of benthic amphipods.

## Introduction

The Southern Ocean (SO) harbours a huge diversity of pelagic and benthic organisms (David & Saucède, 2015; Griffiths, 2010). The long geographic history and isolation of the SO has led to *in situ* evolution of its marine fauna, with adaptations to the cold environment and high levels of endemism (Clarke, 2008; Poulin et al., 2002). The Antarctic marine fauna is experiencing increasing anthropogenic pressure from the exploitation of marine resources and climate change (Constable et al., 2014; Harley et al., 2006; Schofield et al., 2010). In order to conserve the unique Antarctic biota, it is important to understand how biodiversity affects ecosystem functioning.

On of the most dominant groups within the SO benthic communities are the Amphipoda (Peracarida, Crustacea), which show a broad variation in trophic diversity, habitat, lifestyle and size (Dauby et al., 2001; De Broyer & Jazdzewski, 1993, 1996). The superfamily Lysianassoidea is one of the most abundant amphipod taxa in the SO as part of the Antarctic scavenger guild (De Broyer et al., 2004). These amphipods, because of their abundance, play an important role in the energy fluxes of the Antarctic ecosystem, both as scavengers feeding on organic matter and as prey for numerous other organisms (Cherel et al., 2008; Olaso et al., 2000; Panasiuk et al., 2020). Previous studies found that many species within the Antarctic scavenger guild actually relied on multiple feeding types ranging from suspension feeding to opportunistic predation, and as far as true necrophagy (Dauby et al., 2001; De Broyer et al., 2004). Many of the amphipods within the guild have a wide dietary spectrum, thus potentially having high trophic plasticity. However, despite their relevance in the Antarctic marine food web, their eco-functional roles remain poorly understood.

Integrating diversity within trophic levels (horizontal diversity; i.e. taxonomic richness (Ives et al., 2005)) and between trophic levels (vertical diversity; i.e. food chain length and omnivory (Borer et al., 2005)) is one approach to understand the trophic spectra of an ecosystem (Duffy et al., 2007). Above the level of true herbivory (i.e. predators, scavengers), food webs are referred to as tangled with omnivores who tend to feed opportunistically, yet display specificity in their trophic niches (Chikaraishi et al., 2014; Rakusa-Suszczewski et al., 2010; Thompson et al., 2007). Key to understanding the functional diversity and stability of Antarctic food webs, mainly of the scavenger guild, is to attempt to unravel the various bottom-up supplies in these tangled food webs (Bruno & O’Connor, 2005; Chikaraishi et al., 2014).

Stable Isotope (SI) analysis is frequently used to investigate the long-term feeding ecology of organisms in Antarctic food webs (Guerreiro et al., 2015; Michel et al., 2019; Nyssen et al., 2005; Stowasser et al., 2012; Wada et al., 1987). There is a close relationship between the stable isotope ratios assimilated in an organism and its diet (DeNiro & Epstein, 1978; DeNiro & Epstein, 1980). Since primary food sources (i.e. phytoplankton, phytobenthos, and sea ice algae) may vary in stable isotope composition (Wing et al., 2018; Zenteno et al., 2019), stable isotopes of carbon (^13^C:^12^C; δ^13^C) are used to determine the source of primary carbon in food webs. The stable isotope ratio of nitrogen (^15^N^14^N; δ^15^N) is generally measured to determine nitrogen sources and to assess the trophic position of organisms within the food web (Post, 2002). Due to fractionation of the isotopes, consumers are generally enriched in ^15^N through their diet, resulting in a sharp increase of δ^15^N values with each trophic level (DeNiro & Epstein, 1980; Nienstedt & Poehling, 2004). Combining C and N isotope ratios allows to compare the isotopic niche (i.e. a proxy of trophic niche) of different species (Jackson et al., 2011; Newsome et al., 2007).

Our aim is to decipher the trophic niche of *Charcotia obesa* and *C. amundseni*, previously known as the single species *Waldeckia obesa* (Chevreux, 1906). The two sister species were recently described by d’Udekem d’Acoz et al., (2018) based on morphological, molecular and bathymetric data. *Charcotia obesa* occurs in a depth range from 0-150 m (with decreasing occurrences below 120 m), whereas *C. amundseni* is present from 120 to 1000 m depth, resulting in a narrow overlap in depth distribution. Previous dietary studies on *W. obesa* demonstrated a scavenging lifestyle and resistance to starvation (Chapelle et al., 1994; Dauby et al., 2001; Janecki & Rakusa-Suszczewski, 2005). Lysianassoid species in the SO vary strongly in their feeding ecology, even between closely related species (Havermans et al., 2010; Seefeldt et al., 2018). The morphological differences found by d’Udekem d’Acoz et al. (2018) include some variations in the upper lip complex between both species. Nevertheless, the differences are relatively small and, moreover, the morphology of the feeding appendages cannot conclusively deduce the feeding type and the trophic niche of these species (Dauby et al., 2001; Michel et al., 2020). Hence, we hypothesize that, despite small differences of the mouth parts, there is no difference in trophic niche between the two species. Additionally, we expect spatial variation within the two amphipod species in trophic ecology among sampling stations. Since *Charcotia* amphipods are scavenging omnivores, likely to feed opportunistically on dead organic matter, will therefore display the strong seasonal variation of primary production in the SO in their stable isotope ratios (Nygård et al., 2012).

## Materials and methods

### Sample collection

Specimens from the genus *Charcotia* were obtained from collections curated at various scientific institutes (i.e. the British Antarctic Survey, University of Lodz, Alfred Wegener Institute, Muséum National d’Histoire Naturelle Paris and the Royal Belgian Institute of Natural Sciences). Specimens were collected from various locations and depths in the SO (Table 1) during several scientific expeditions. Amphipods were caught using epibenthic sledge or baited traps, sorted, identified and preserved in precooled molecular grade ethanol (96-99 %) at −20°C. Samples originated from four geographic areas (Figure 1): the Weddell Sea near the Filchner Area, the West Antarctic Peninsula (WAP), Queen Maud Land and the Adélie Coast. *Charcotia obesa* Chevreux, 1906 (Crustacea, Amphipoda, Lysianassoidea) was sampled in seven locations, six off the south-west shelf of the Antarctic continent (Admiralty Bay (AdB); South Shetland Island (SSI); Foyn Harbor (FH); Palmer Island (PAL); Andvord (AND); Berthelot Islands (BI)) and one on the south-east coast (Dumont d’Urville Sea (URV)). Samples in the south-west are subcategorized as two locations in the north (AdB, SSI), three central (FH, PAL, AND) and one in the south (BI). *Charcotia amundseni* d’Udekem d’Acoz 2018 (Crustacea, Amphipoda, Lysianassoidea) was collected at two geographical locations, one in the Weddell Sea (Filchner Area (FIL)) and two off the coast of Queen Maud Land (Breid Bay (BB); Crown Bay (CB)). The final dataset contained 205 *C. obesa* individuals and 41 *C. amundseni* individuals, based on morphological identification and confirmation with mitochondrial DNA COI sequences.

**Table 1.**
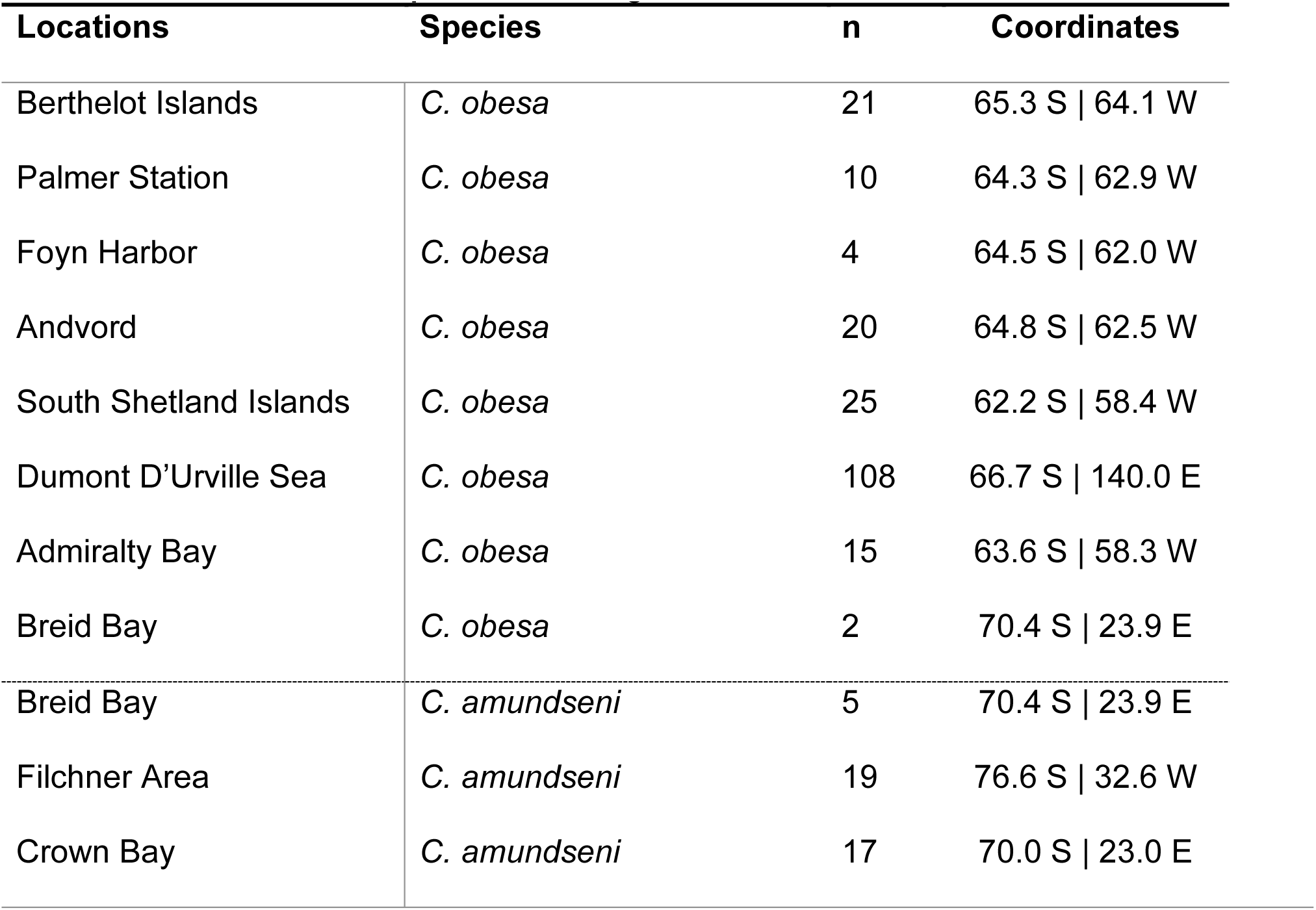
Sampling stations of *Charcotia obesa* and *C. amundseni*, including the number of individuals sampled at each station and the coordinates. For the exact geographic locations around the Antarctic continent, please refer to Figure 1.

**Figure 1.**
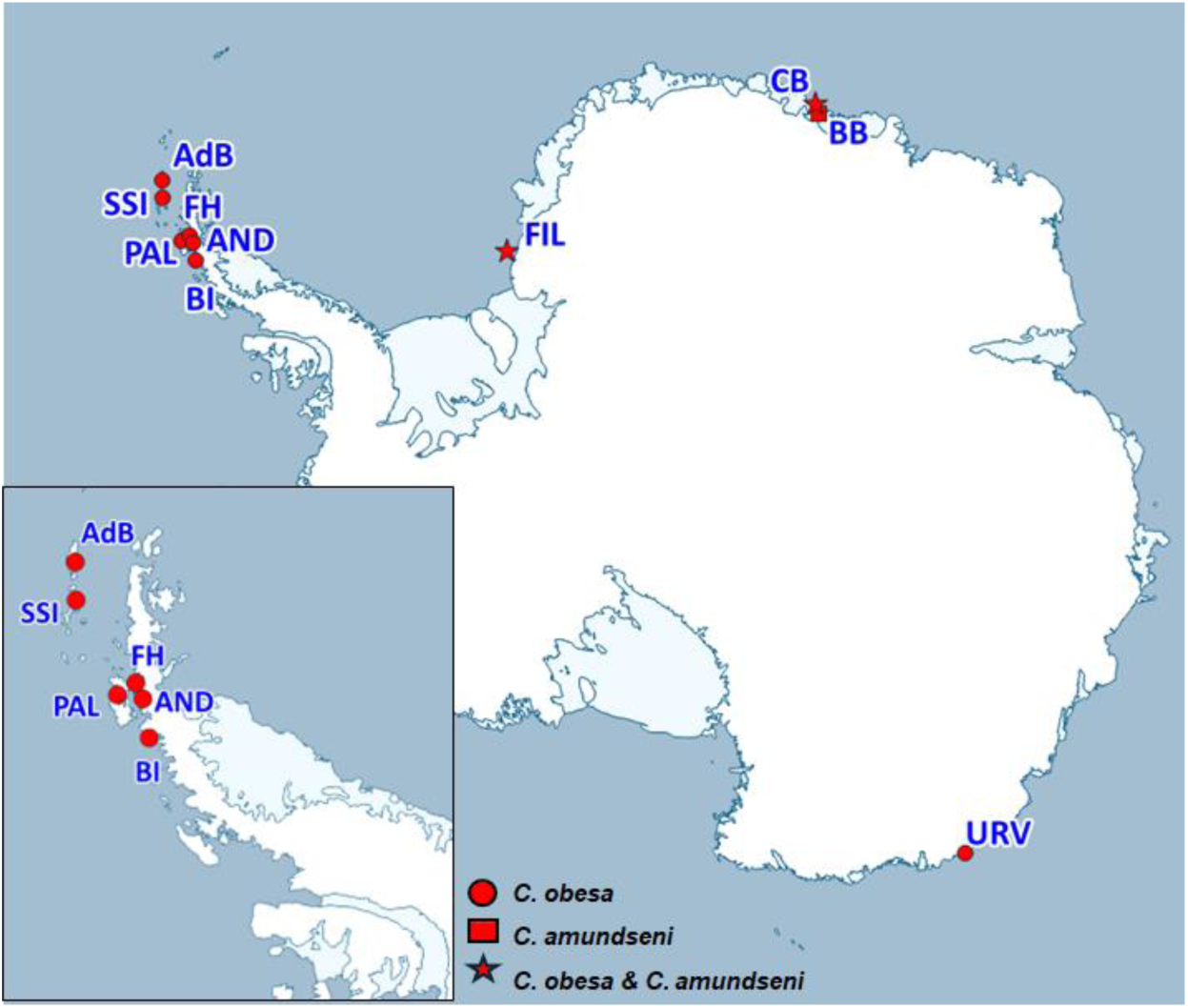
Sampling locations of Charcotia obesa (circles) and C. amundseni (squares) as well as sampling locations with both (stars). This map was made with the QGIS (Version 3.16, QGIS.org) add-on Quantarctica (Version 3.2, Matsuoka et al., 2018).

### Stable Isotope Analysis

Two to four pleopods (depending on size) including muscles were dissected from each *Charcotia* specimen and air-dried for at least 72 h. By using the same tissue (i.e. mostly soft tissue muscles) for all samples, we exclude potential bias and underestimations due to the analysis of different tissue (Søreide & Nygård, 2012). The protocol allows to conserve collection specimens for taxonomic purpose and future studies. Pleopods were weighted in tin cups; dry weight ranged from 0.1 to 0.4 mg. Samples were analysed at University of Liège, using an Isotope Ratio Mass Spectrometer (IRMS) (precisION, Elementar) coupled to an elemental analyser (VarioMicrot, Elementar). Delta (δ) notations of carbon (δ^13^C) and nitrogen (δ^15^N) were used to express isotope ratios, which are calculated here as parts per thousand (‰) (Coplen, 2011). We used blank tin cups, secondary analytical material (glycine and European sea bass *Dicentrarchus labrax* reference material) and certified material from the International Atomic Energy Agency (IAEA, Vienna, Austria), IAEA C-6 (sucrose; δ^13^C = −10.8± 0.5 ‰) and IAEA-N1 (ammonium sulphate; δ^15^N = 0.4 ± 0.2‰). The isotope ratios are expressed as mean values ± SD for each species and calculated according to sampling station and depth.

### Data analysis

Before SI data was analysed, a correction was applied to account for differences in SI values at different geographical locations. This correction was taken from Le Bourg et al. (2021) and applied to remove the impact of locality (Equation 1).

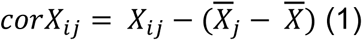

Where *X* is the variable (i.e. δ^13^C), *i* denotes the individual and *j* the sampling station. *X_ij_* describes a value belonging to individual *i* at station *j*, *X̅* is the overall mean of all values and *X̅_j_* is the mean value at station *j*. Accordingly, transformed data are displayed as δ^13^C_corr_ and δ^15^N_corr_. The same formula was used when comparing values from different depth ranges to correct again for possible local effects.

Isotopic niches and overlaps were determined by calculating Standard Ellipse Areas (SEA) to the δ^13^C and δ^15^N patterns, which were corrected for sample-size (SEA_C_), using the SIBER package (V2.1.9, Jackson et al., 2011) in Rstudio v4.3.2 (R Core Team, 2014; RStudio Team, 2020). SEA_C_ were determined for each *Charcotia* species based on all samples, and on samples from the same geographical locations or depths. Additionally, Bayesian estimates of SEA_B_ (based on 2 · 10^6^ iterations, 2 chains, 10^4^ initial discards and a thinning interval of 10) were conducted and compared to SEA_C_. Plots were made with the R package ggplot2 (V3.5.1, Wickham, 2016). When assessing differences in SI ratios at various depths, recorded sample depths were clustered in increments of 50 m, creating four groupings of *C. obesa* samples at depths of <50 m (n = 115), 50 to 100 m (n = 29), 100 to 150 m (n = 34) and >150 m (n = 25) (Table 5). *Charcotia amundseni* samples were not considered for this type of analysis as all samples were within 50 m from each other, at a depth range from 230 to 274 m. To estimate the trophic position (TP) of both species, the tRophic position package (V0.8.0, Quezada-Romegialli et al., 2018) was used. To establish a baseline, we used collated data on Particulate Organic Matter (POM) from an earlier study (St John Glew et al., 2021) and compared the uncorrected data to the Antarctic pelagic baseline. TP is noted as the statistical mode with a 95 % Credibility Interval (CI_95_).

For statistical analysis, data outliers were detected and removed using the Rosner test; this resulted in the removal of three samples from *C. obesa*. Normality was tested with the Shapiro-Wilkes’ test and homoscedasticity with the Levene’s test. Since parametric assumptions were not met, Mann Whitney U tests, Kruskal-Wallis tests and Chi-squared tests were used. *Post hoc* testing was performed with Dunn tests (Bonferroni corrected). A significance level of p-value < 0.05 was used in all tests.

## Results

### Comparing SI ratios between two

Charcotia *species.* δ^13^C ranged from −20.3 ‰ to −25.6 ‰ and δ^15^N varied from 8.2 ‰ to 13.6 ‰ among the 205 samples of *C. obesa*. The total dataset of 41 *C. amundseni* samples resulted in δ^13^C and δ^15^N ranging from −20.8 ‰ to −24.3 ‰ and from 11.1 ‰ to 15.0 ‰, respectively (Figure S1). Having corrected for variation of SI ratios between locations as explained in Materials and Methods, *C. amundseni* display a higher mean δ^15^N_corr_ of 12.9 ± 0.7 ‰ (Min. 10.9 – Max. 14.4 ‰) compared to the δ^15^N_corr_ of *C. obesa* of 10.7 ± 0.6 ‰ (Min. 9.1 – Max. 12.2 ‰) (Figure 2; Table 2). *Charcotia amundseni* showed a significantly higher δ^15^N_corr_ than *C. obesa* (W = 7712, p-value < 0.001) (Table 3). In contrast, δ^13^C_corr_ values for both species were rather similar, with *C*. *amundseni* having a mean δ^13^C_corr_ of −23.7 ± 0.5 ‰ (Min. −24.2 – Max. −21.1 ‰) and *C. obesa’s* mean δ^13^C_corr_ of −23.8 ± 0.4 ‰ (Min. −25.2 – Max. −21.8 ‰) (Figure 2; Table 2). The δ^13^C_corr_ values of *C. amundseni* and *C. obesa* did not show any significant difference (W = 5008, p-value = 0.053) (Table 3). Ellipse-based metrics computed using the SIBER package showed no measurable overlap between both *Charcotia* species (Fig. 2; Table 2).

**Figure 2.**
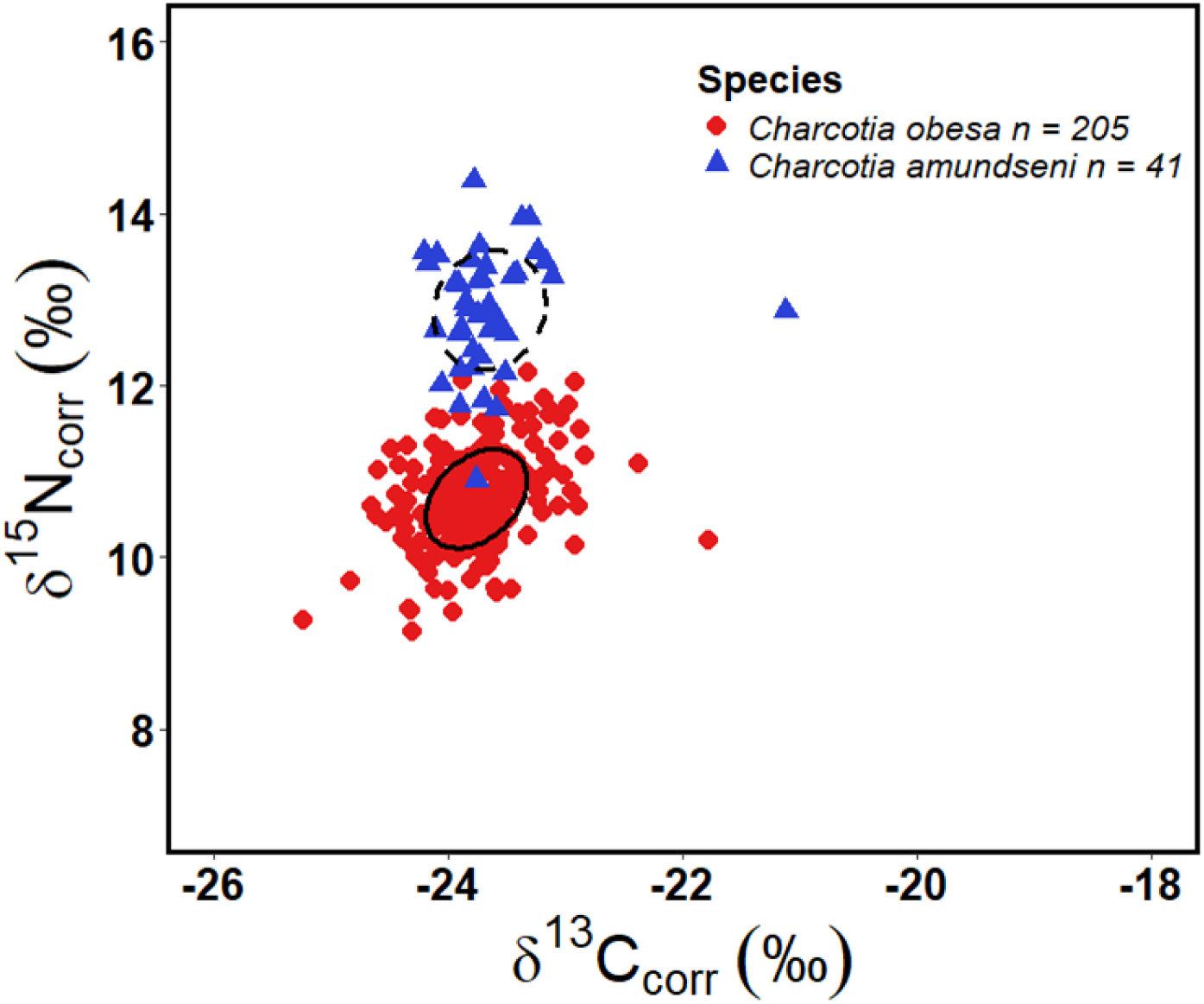
Standard Ellipse Areas of individual stable isotope values of δ13C and δ15N, corrected for locations, and its isotopic niche, of Charcotia obesa (red circles, solid ellipse) and C. amundseni (blue triangles, dashed ellipse).

**Table 2.** Mean ± SD δ^13^C and δ^15^N values calculated *for Charcotia obesa* and *C. amundseni*. Corrected values (corr) are adjusted values of δ^13^C and δ^15^N for their respective locations.

| Species | n | $\delta^{13}\text{C}$ (‰) | $\delta^{15}\text{N}$ (‰) | $\delta^{13}\text{C}_{\text{corr}}$ (‰) | $\delta^{15}\text{N}_{\text{corr}}$ (‰) | Overlap |
| --- | --- | --- | --- | --- | --- | --- |
| *C. obesa* | 205 | -23.7±1.0 | 10.7±1.2 | -23.8±0.4 | 10.7±0.6 | <0.01 % |
| *C. amundseni* | 41 | -23.6±0.6 | 12.9±0.9 | -23.7±0.5 | 12.9±0.7 | |

**Table 3.**
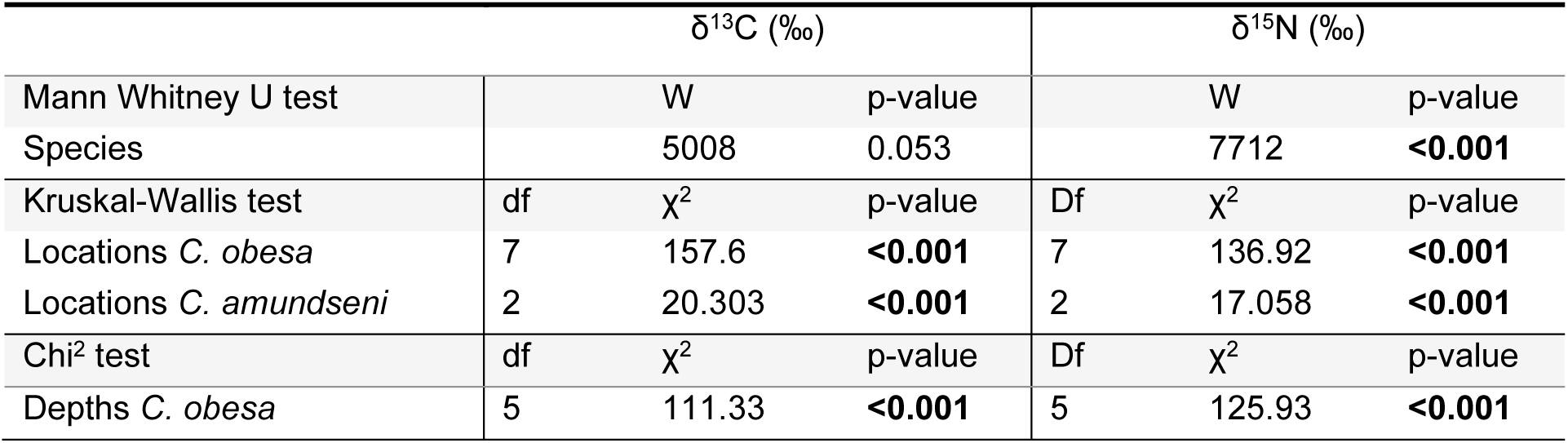
Results of pairwise t-test, ANOVAs and Kruskal-Wallis tests assessing the influence of species, locations and depths on stable isotope values of carbon and nitrogen. Significant p values (<0.05) are indicated in bold.

Finally, using the uncorrected δ^15^N values from both species, we calculated a trophic position of 4.19 (CI_95_: 3.66 – 4.97) for *C. obesa* (Figure 3), and 5.14 (CI_95_: 4.46 – 6.17) for *C. amundseni*. Our trophic position model suggested that this difference was meaningful, with the probability of *C. amundseni* occupying a higher trophic position being 96.1%.

**Figure 3.**
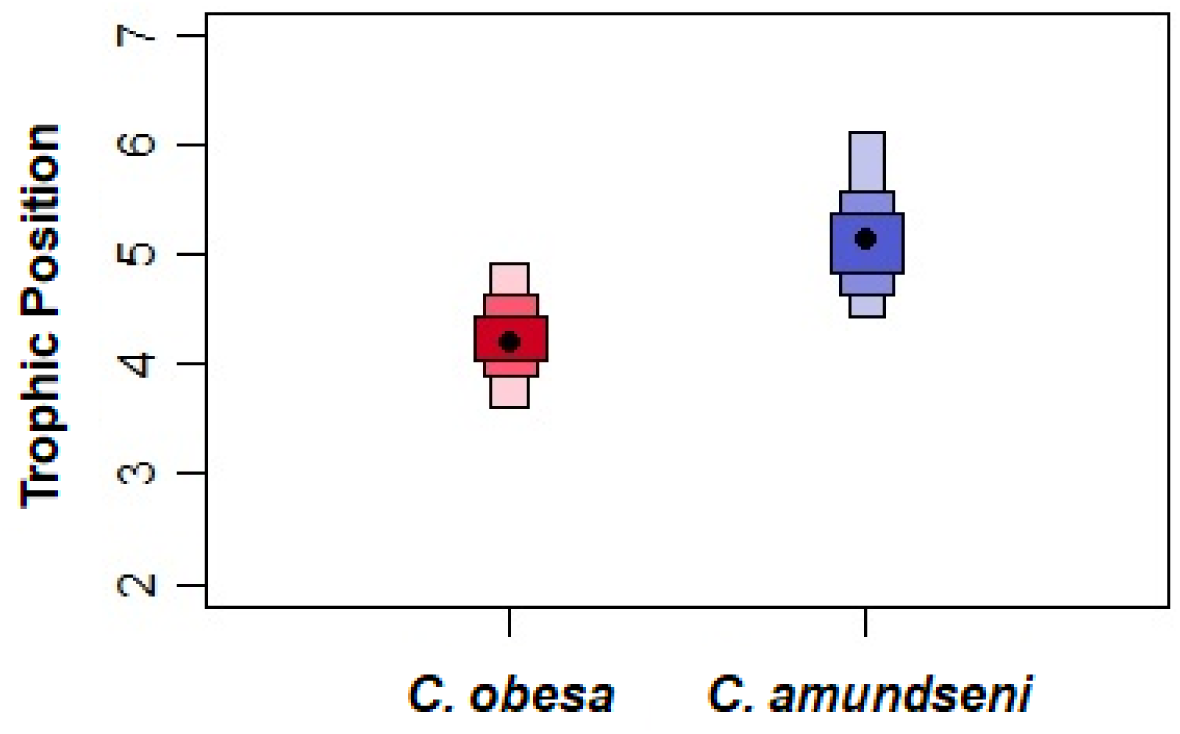
Trophic positions calculated for Charcotia obesa (red) and C. amundseni (blue). Boxplots show credibility intervals of 50, 75 and 95 %. Black dots denote the mode of each species’ trophic position.

### Comparing SI ratios between geographic locations

This study shows that there were significant differences in SI ratios of carbon and nitrogen for both *Charcotia obesa* and *C. amundseni* between the sampled regions of the Antarctic continental shelf (Table 4). The lowest δ^13^C was observed in both Palmer Station (PAL) and the South Shetland Islands (SSI) (−24.9 ± 0.3 ‰), while the highest δ^13^C ratios were observed in Andvord (AND) (−22.0 ± 0.6 ‰). The lowest value of δ^15^N is reported in Admiralty Bay (AdB) (9.2 ± 0.8 ‰) and the highest δ^15^N is observed at the Berthelot Islands (BI) (12.6 ± 0.6 ‰). Carbon SI ratios of *C. obesa* were significantly different in 14 out of 21 pairwise difference tests at seven sampling locations. Nitrogen SI ratios were significantly different in 12 out of 21 pairwise tests.

**Table 4.** Sample size (n) and mean ± SD δ13C and δ15N values calculated for *Charcotia obesa* and *C. amundseni*; values are grouped per sampling location.

| Species | Locations | n | $\delta^{13}\text{C}$ (‰) | $\delta^{15}\text{N}$ (‰) |
| --- | --- | --- | --- | --- |
| *C. obesa* | Admiralty Bay | 15 | -24.7±0.5 | 9.2±0.8 |
| | South Shetland Islands | 25 | -24.9±0.3 | 9.8±0.5 |
| | Palmer Station | 10 | -24.9±0.3 | 11.6±0.3 |
| | Foyn Harbor | 4 | -23.3±0.1 | 12.1±0.6 |
| | Andvord | 20 | -22.0±0.6 | 12.2±0.6 |
| | Berthelot Islands | 21 | -22.2±0.7 | 12.6±0.6 |
| | Dumont D'Urville Sea | 108 | -23.9±0.4 | 10.3±0.6 |
| *C. amundseni* | Breid Bay | 5 | -24.1±0.2 | 11.8±0.7 |
| | Crown Bay | 17 | -23.9±0.2 | 12.6±0.5 |
| | Filchner Area | 19 | -23.4±0.7 | 13.5±0.9 |

When checking for a possible geographic grouping of locations, individuals belonging to the northern group (AdB & SSI) had significantly lower δ^15^N and δ^13^C ratios (p-value <0.001) compared to individuals found in the central (PAL, FH & AND) and especially the southern group (BI) (p-value <0.001) (Table 5a). Within-group pairwise comparisons of locations around the WAP of both δ^15^N and δ^13^C, indicated that adjacent stations are non-significant, with the exception of δ^13^C in the central group, (Table 5 b-c) and an increasing gradient from north to south of both δ^15^N and δ^13^C was observed (Table 4). The only sampling station (Dumont d’Urville Sea, URV) in the South-East differed significantly from all other stations in both δ^15^N and δ^13^C except for Foyn Harbor (FH) concerning the δ^13^C and FH and SSI regarding δ^15^N. Standard ellipse overlap was also minimal between most locations (<5 %), except for AND and BI (30.95 %) (See Figure 4a, Figure 5a and Table 5d).

**Table 5.**
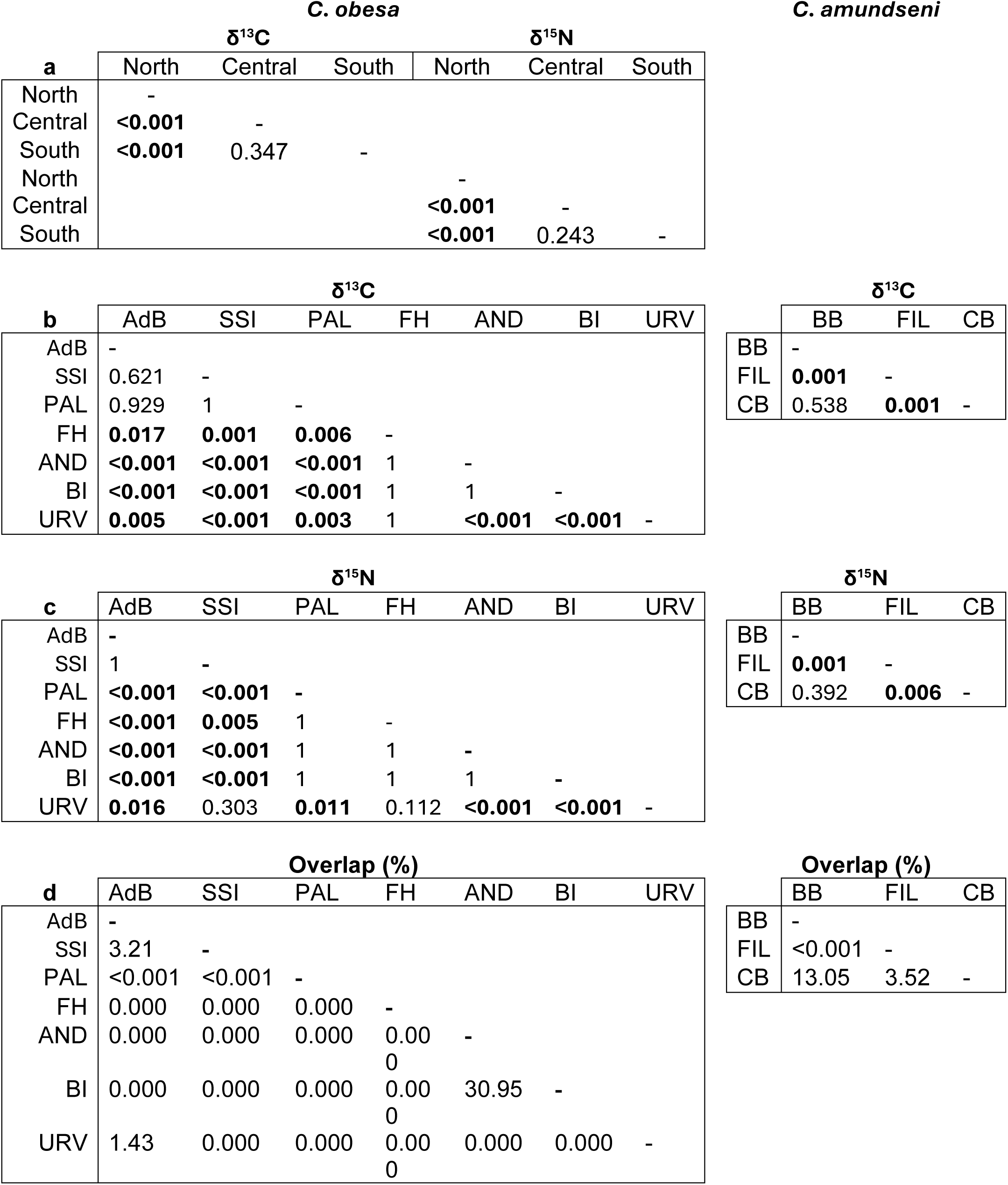
Results of pair-wise Tukey honest significance tests assessing the influence of location on stable isotope values. (a) pair-wise tests between the North, Central and South clusters of locations. (b) pair-wise tests between locations for δ13C for both *Charcotia* species. (c) pair-wise tests between locations for δ15N for both *Charcotia* species. Significant p values (<0.05) are indicated in bold. (d) overlap between geographical locations, displayed in percentages.

**Figure 4.**
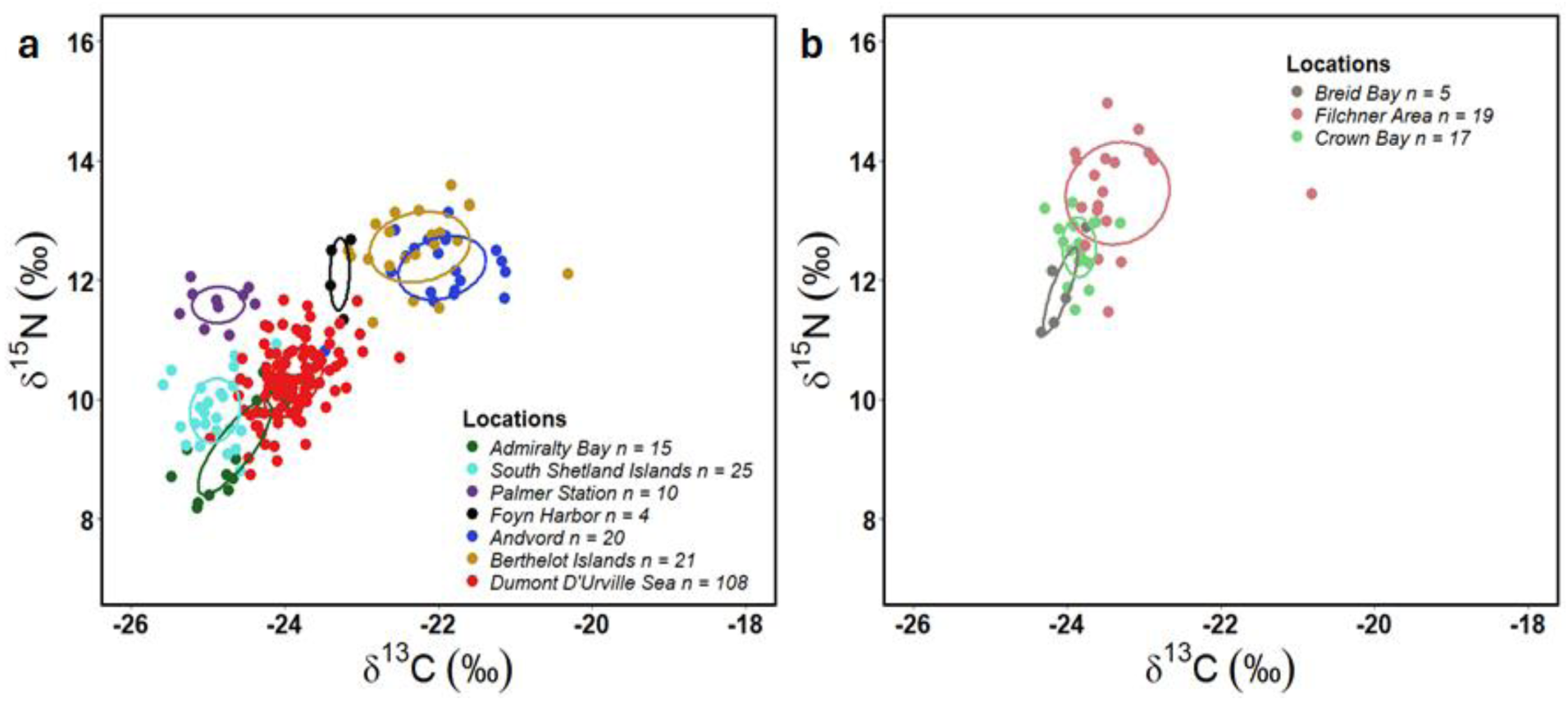
Standard Ellipse Areas of individual stable isotope values of δ13C and δ15N for Charcotia obesa (a), C. amundseni (b) and their isotopic niches, grouped per location. Locations are indicated by different colours; the sample number per location is provided in the legend.

**Figure 5.**
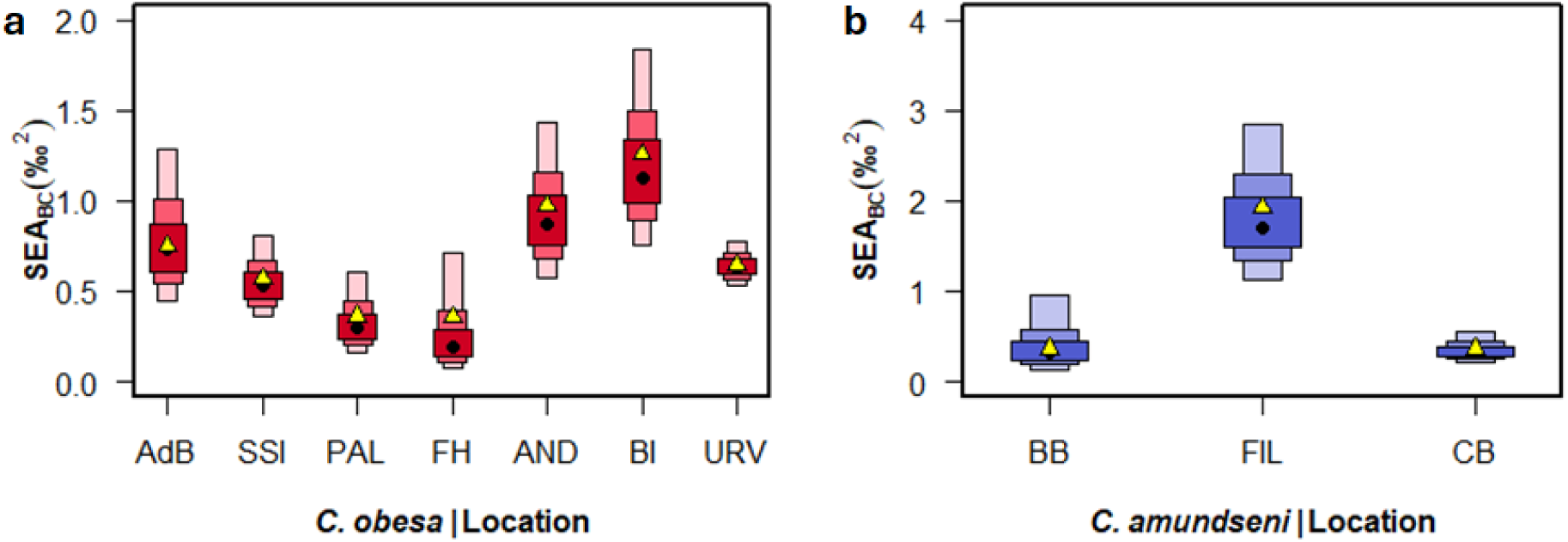
Standard Ellipse Areas(BC) of stable isotope values of geographical locations for both Charcotia obesa (a, red) and C. amundseni (b, blue). Black dots denote the Bayesian estimate of SEA while yellow triangles indicate the computed SEA.

Three locations were sampled for *C. amundseni*. Both Crown (CB) and Breid Bay (BB) are geographically adjacent at Queen Maud Land. The lowest δ^13^C were found in BB (−24.1 ± 0.2 ‰) while the highest δ^13^C was in the Filchner Area (FIL) (−23.4 ± 0.7 ‰). Similarly, the lowest δ^15^N was reported from BB (11.8 ± 0.7 ‰) while the highest δ^15^N is found in the FIL (13.5 ± 0.9 ‰) (Table 4; Figure 4b). δ^13^C differed significantly between specimens from the FIL (n = 19) compared to CB (n = 17) (p-value = 0.001) and BB (n = 5) (p-value = 0.001), same patterns are shown in δ^15^N with significant differences between BB (n = 5) and the FIL (n = 19) (p-value < 0.001) and between the FIL (n = 19) and CB (n = 17) (p-value = 0.006). Overall, the location within Weddell Sea (FIL) and two locations of Queen Maud Land (CB and BB) showed significant differences in both δ^15^N and δ^13^C, whereas the variation between the two adjacent sites is much lower (Table 5 b-c). Niche overlap was determined for all locations using standard ellipses (Figure 5b), showing that CB had slight overlap with both BB (13.05 %) and the FIL (3.52 %), but BB and FIL had no overlap between each other (Table 5d).

### Depth-dependent SI ratios

*Charcotia obesa* displayed a large overlap in isotopic niche between depths (Figure 2S/Table 1S). Moreover, values for both δ^13^C_corr_ and δ^15^N_corr_ differed only a few decimal points (Table 2S).

## Discussion

The isotopic composition of carbon (δ^13^C) and nitrogen (δ^15^N) of two recently described *Charcotia* sister species (*C. obesa* and *C. amundseni*, d’Udekem d’Acoz et al., 2018) were analysed to compare their trophic position in the Antarctic food web. Significant differences in nitrogen isotope ratios were found between the two sister species. Based on the high nitrogen values, high trophic levels were estimated for both species. Significant differences in both stable isotopes between regions were found within each species. An increasing trend in both stable isotopes in *C. obesa* was found from the north to south off the WAP. Significant differences were also observed in *C. amundseni* from sampling locations in the Weddell Sea and Queen Maud Land.

### Nitrogen SI ratios and trophic positions

δ^15^N values for both species were high, corroborating a high position in the trophic web and, probably, a necrophagous scavenging behaviour (Dauby et al., 2001; Nygård et al., 2012). *Waldeckia obesa*, a mix of *C. obesa* and *C. amundseni*, showed δ^15^N values of 11.6 ± 0.3 ‰ in the eastern Weddell Sea (Nyssen et al., 2002) and lower values along the Antarctic Peninsula (7.3 ± 0.7 ‰) (Nyssen et al., 2005). Michel et al., (2019) have reported values of 9.1 ± 1.6 ‰ in East Antarctica (Adélie Land), which differs slightly with the isotopic ratio of *C. obesa* in our study (Table 4). δ^15^N ratios in the previous studies match the range found in our study (Min. 9.1 – Max. 12.2 ‰). Other scavenging amphipods investigated by Zenteno et al. (2019) had δ^15^N values ranging from 4.4 ± 0.6 ‰ to 5.9 ± 0.3 ‰, with the highest δ^15^N value (6.6 ± 0.4 ‰) for a carnivore predatory amphipod. One species included in the latter study was *Cheirimedon femoratus* Pfeffer, 1888 (Crustacea, Amphipoda, Lysianassoidea), which is considered an omnivorous scavenger from the same scavenging guild as *Charcotia* (De Broyer et al., 2004; Seefeldt et al., 2017, 2018)*. Cheirimedon femoratus* occupies a broad trophic range within the guild (Seefeldt et al., 2017). This species is notably smaller than *Charcotia* and has a different mandible morphology. *Charcotia* feeds on carcasses from outside to inside, while smaller species obtain access either through orifices in the body or from openings created by larger scavengers, such as *Charcotia* (Seefeldt et al., 2017). *Charcotia amundseni* and *C. obesa* displayed different δ^15^N_corr_ values, suggesting a different trophic position (TP; Bearhop et al., 2004; Layman et al., 2012; Le Bourg et al., 2020; Post, 2002). *C. amundseni* (TP of 5.14) presumably belonged to the fifth trophic level and *C. obesa*, with an estimated TP of 4.19, to the fourth trophic level. A first explanation could be that, although the two species are opportunistic scavengers, *C. amundseni* which is found deeper feeds on carcasses occupying a higher trophic position than *C. obesa*. *Charcotia* are feeding episodically and are often found on large dead animals at depth (Bolstad et al., 2023; Jażdżewska, 2009). We hypothesize that in shallow areas, the diversity of dead prey is larger and belongs to more diverse trophic positions than in deeper areas. Deep-sea carcasses are typically dominated by larger organic falls of organisms from higher trophic levels (Bolstad et al., 2023). Consequently, explaining the lower trophic position of *C. obesa* compared to *C. amundseni*. Anyway, our modelled results show that the two species occupy very high trophic positions, much higher than the one observed by Michel et al. (2019) (*C. obesa*, TP: 2.4) for example. This difference is huge in terms of energy flow and could indicate that *C. obesa* has higher feeding plasticity then initially thought. In East Antarctic, *C. obesa* was caught at shallow depths (20 m), preying likely on dead invertebrates occupying a low trophic position. Our *Charcotia* species are at a higher trophic position than smaller scavenging amphipods investigated by Zenado et al. (2019), showing that the scavengers guild does not necessarily depend on the same prey and comprises organisms occupying different trophic positions in the Antarctic food web (Smale et al., 2007). Niche partitioning might allow scavenging amphipods to co-exist in the same habitat.

Differences in trophic position and thus in effective ecological niche support the coexistence of morphologically similar species and contribute to the diversification of sister species (Bessa et al., 2014; Klarner et al., 2013). Nevertheless, a second hypothesis could be that both species display eco-physiological differences. Isotopic fractionation is associated with starvation, leading generally to an increase of δ^15^N values (Doi et al., 2017). Impacts of diet quality and starvation on nitrogen isotopic fractionation have been reported in other arthropods (Haubert et al., 2005) and crustaceans (Fantle et al., 1999; Trochine et al., 2019). Starvation has been reported for *W. obesa* (Chapelle et al., 1994), which is an opportunistic omnivorous scavenger eating episodically. The higher δ^15^N values observed in *C. amundseni* as opposed to *C. obesa*, might indicate that *C. amundseni*, living in deeper water, cope with starvation more frequently and for elongated periods of time than the shallow water *C. obesa* individuals. Additionally, recent research found that differences in gut microbiome composition and specific symbiotic relationships aid deep sea invertebrates to cope with surviving in these unusual environments. Microbiome diversity leads to variable host physiology, behaviour and ecology (Osman & Weinnig, 2021) and is alternative explanation for the trophic differences found in this study.

### Carbon SI ratios and food sources

No significant difference was found between the mean δ^13^C_cor_ values of both species (Table 3). δ^13^C values are generally indicative of the primary food source of an organism (France, 1995; Michener & Kaufman, 2007) with more negative δ^13^C values (± −25 ‰ to ± −30 ‰) being characteristic of pelagic primary producers (Espinasse et al., 2019; Michel et al., 2019), intermediate δ^13^C values (± −25‰ to ± −10‰) of benthic primary producers and less negative δ^13^C values (± −20 ‰ to ± −8 ‰) characteristic of sea ice microbial communities (Gillies, et al., 2012; Gillies et al., 2013; Michel et al., 2019). *Charcotia obesa* has the largest range in δ^13^C, which might be attributed to the larger sample size, but also reflects a wide diversity of primary food sources (both pelagic and benthic) and larger diversity in dead organic matter in the shallower depth range in which they occur. The range of δ^13^C values of *C. amundseni* falls within the predetermined range of benthic primary producers as primary food source (Gillies et al., 2012; Michel et al., 2019). A wide range in δ^13^C values indicates the use of different primary food sources, confirming their scavenging foraging feeding behaviour (Amsler et al., 2014; Aumack et al., 2017). The study by Zenteno et al. (2019) found δ^13^C values for scavenging amphipods ranging from −14.7 ± 0.6 to −21.5 ± 0.6 ‰, aligning with our results. The availability of multiple primary food sources is one of the key features that enables the stability of Antarctic food webs (Zenteno et al., 2019).

### Effect of location

The primary food sources in Antarctic food webs (phytoplankton, macroalgae, and detrital matter) are highly dependent on sea-ice coverage, and therefore also shaped by location and seasonality (Norkko et al., 2007). Our results confirm this, as there is limited overlap between sites on the δ^13^C -axis (see Figure 4a). When investigating the trophic ecology of each species, carbon isotope ratios of *C. obesa* individuals showed significant geographical differences between localities off the Antarctic continent (Table 3 and 5 b-c). Lowest mean δ^13^C_cor_ values (−24.8 ‰) were estimated around the SSI while the highest mean δ^13^C_cor_ value (−22.0 ‰) was measured in Andvord (AND) (Table 4). Interestingly, Dumont d’Urville Sea (the only south-east sampling location) differed significantly in the mean δ^13^C_cor_ value from all other locations except Foyn Harbor (with the lowest sample size per location). The biogeochemistry of the Southern Ocean differs regionally, which affects primary production (Fraser et al., 2023; Henley et al., 2020) and is reflected in our isotope data. For *C. amundseni*, samples from three locations were included in the current study and we found a significant difference between the Weddell Sea location (Filcher Area (FIL)) and both locations in Queen Maud Land (Breid Bay (BB); Crown Bay (CB)) (Table 5b). The carbon stable isotope ratio is highly influenced by sea surface temperature and CO_2_ availability for photosynthesis (Espinasse et al., 2019; Lara et al., 2010). Additionally, primary production is negatively correlated with sea ice coverage, leading to an increase of detritus consumption by benthic consumers, which is reflected in higher δ^13^C values (Norkko et al., 2007). δ^15^N values, on the other hand, are strongly influenced by sea ice dynamics and upwelling, since the availability of nitrogen (in the form of nitrate) is the limiting factor and will cause an increase in stable isotope values (Difiore et al., 2010; Zenteno et al., 2019). The lowest mean value of δ^15^N_cor_ (9.0 ‰) in *C. obesa* was estimated in Admiralty Bay (AdB), while the highest mean value (12.8 ‰) was found in Berthelot Islands (BI), showing an increasing gradient of 3 ‰ from the northernmost sampling location around the WAP to the most southwest sampling location close to the Bellingshausen Sea. A similar pattern was found by Brault et al. (2018) in zooplankton species from the WAP to the Ross Sea. The authors attributed this pattern to the abundant polynyas and higher productivity due to glacial inputs of iron in the Amundsen and Ross Sea, leading to a more δ^15^N enriched phytoplankton (Brault et al., 2018). Similar meltwater input has been reported in the Bellingshausen Sea, which could explain the observed pattern of increasing δ^15^N (Holland et al., 2010; Sheehan et al., 2023) in our study. Significant differences in nitrogen stable isotopes of *C. amundseni* were found between FIL and both other locations (CB and BB). The Filchner Area is located on the east side of the Weddell Sea, adjacent to the Weddell Gyre which contributes to upwelling of nutrient rich deep-sea water. The latter might be linked to the higher δ^15^N values (Gordon et al., 2001; Nicholls et al., 2009; Vernet et al., 2019).

### Depth effect

*Charcotia obesa* samples originated from four depth ranges along the shelf of the Antarctic continent (see Tables S1 & S2, Figure S2). Distance to the continent and consequently also depth can have a strong influence on both isotopic ratios. Usually, coastal environments show higher δ^13^C and δ^15^N ratios, which decrease with distance to the shore (Lara et al., 2010; Zhang et al., 2014). The decrease in the carbon isotope ratio is generally steeper than for the nitrogen isotope (El-Sabaawi et al., 2012; St John Glew et al., 2021). However, in our data, this decrease is not very pronounced and the four depth ranges show complete overlap in the SEA analysis (Figure S2).

### Suggestions for future studies

Our study illustrates that a limited number of geographical stations can produce valuable novel insights. Increasing the sampling size to at least five individuals (and equally distributed) per location and a more structured distribution of locations and depths around the Antarctic continent might make the statistical analyses across variables more robust. Isotopic measurements of POM during sampling would also be advisable to provide baseline values for the species of interest (Michel et al., 2016). Sampling outside of the Austral summer is recommended, although challenging if not next to impossible. Increasing the temporal resolution and year-round sampling could further improve the accuracy of observed patterns (de Lima et al., 2022; Kolts et al., 2013).

Stable isotopes are a powerful tool to estimate trophic niches of organisms but may lack resolution especially when spatiotemporal variation in the ecosystem is high. Including dietary studies by sampling the stomach content might provide a more detailed, albeit snapshot, insight in the diet and starvation periods. Metabarcoding of the stomach content has proven to be a useful tool at a higher resolution than visual dietary assessments (Maes et al., 2022). Therefore, a combination of trophic markers, molecular and morphological methods will result in information at the highest resolution and predict a species’ trophic niche most accurately (Gerringer et al., 2017).

## Conclusion

Based on the data presented here, we observe that even closely related sister species, which occupy similar habitats and have similar feeding strategies differentiate their trophic niche. The plasticity in feeding habits of scavenging amphipods might have important implications in the face of climate change. Polar regions are one of the fastest warming regions in the world, more specifically the Antarctic Peninsula is affected to a great extent (Engel et al., 2024; Wallis et al., 2023). Decreasing sea ice cover might alter the effective niche of certain species and reduce suitable habitat (Parkinson & Cavalieri, 2012). Trophic plasticity of Antarctic benthic organisms could be a strategy to ensure survival, however potentially leading to different responses of each group (Michel et al., 2016). Therefore, future focus on trophodynamics is essential in terms of insights in ecosystem functioning and conservation of the pristine Antarctic environment.

## Acknowledgements

We thank all scientists and staff from Muséum National d’Histoire Naturelle, Paris; British Antarctic Survey, Cambridge; Alfred Wegener Institute, Bremerhaven; University of Lodz, Lodz; Institute of Natural Sciences; Brussels for providing specimens from their collections for the current study. We also thank scientists and crew from the Antarctic expeditions BEL121, PS81, PS129, ANTARXXVVII for collecting specimens. Special thanks to B. Danis who kindly deployed some baited traps during his visit to the Princess Elisabeth Station in 2022 (funded by the RECTO project). F. Volckaert provided valuable comments to the manuscript. This study was funded by BELSPO, call BRAIN.be 2.0 (Contract Nr B2/191/P1/COPE).

## Supplementary

### Figures

**Figure 1S.**
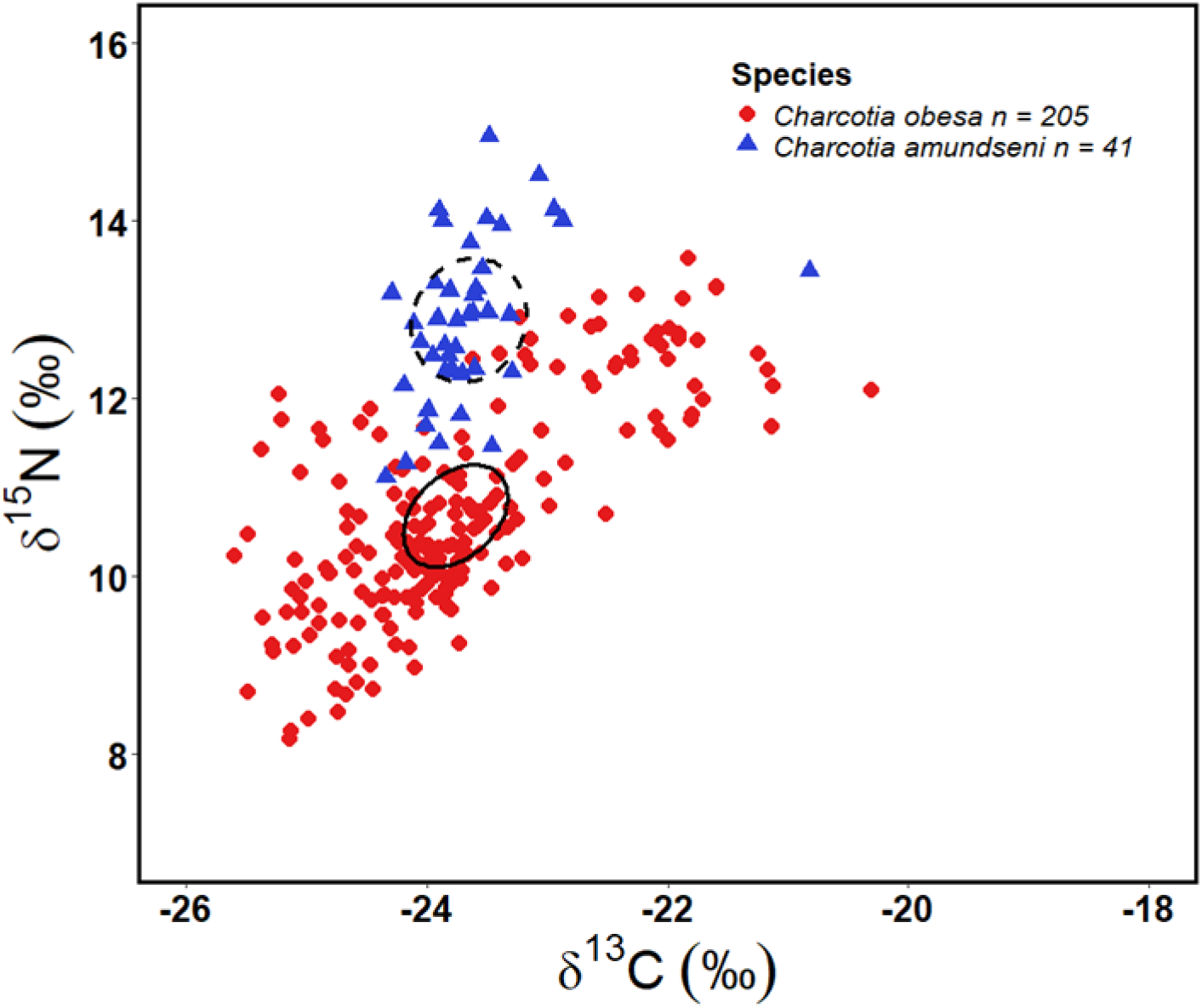
Standard Ellipse Areas of individual uncorrected stable isotope values of δ^13^C and δ^15^N and its isotopic niche, of Charcotia obesa (red circles, solid ellipse) and C. amundseni (blue triangles, dashed ellipse).

**Figure 2S.**
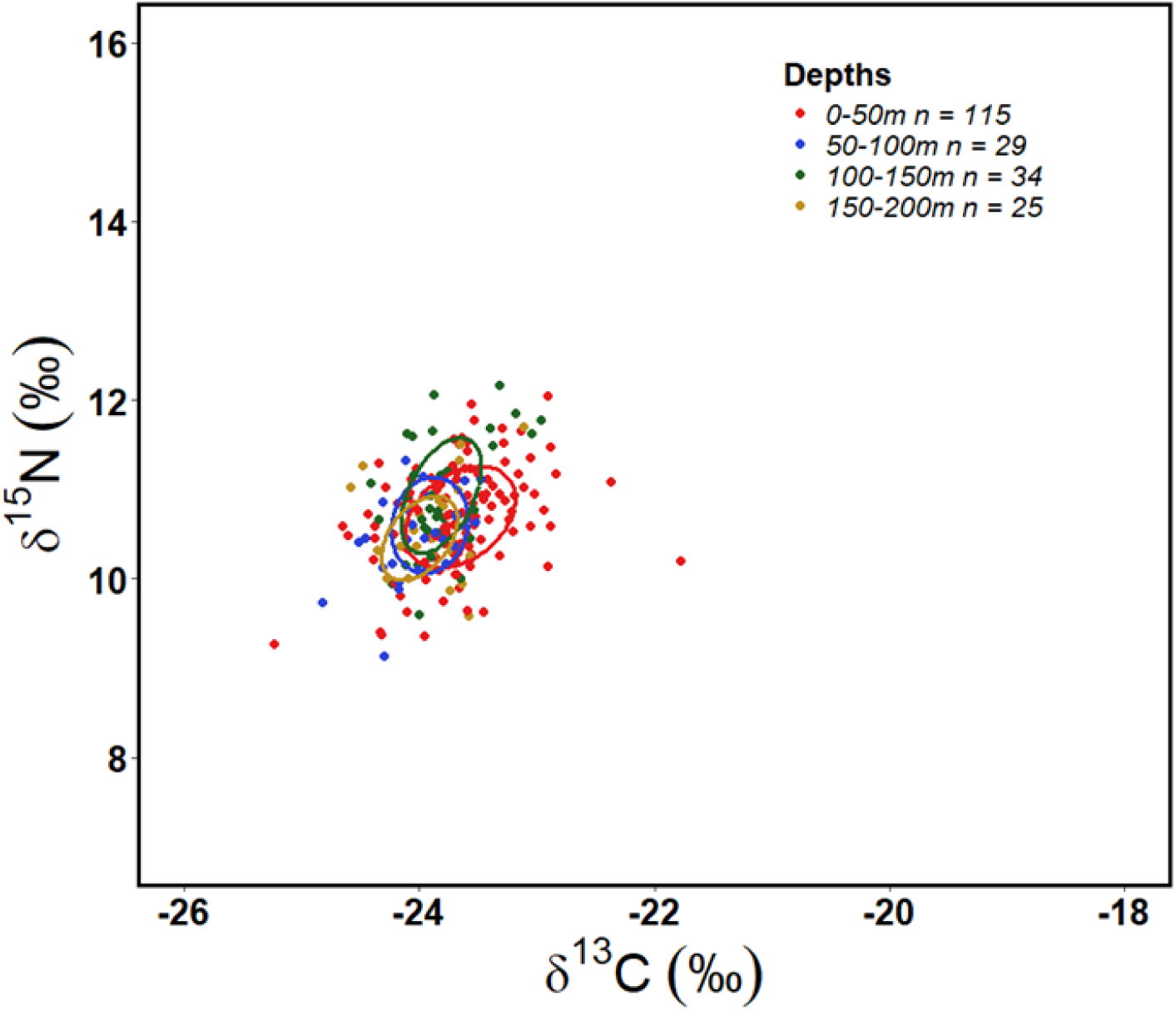
Standard Ellipse Areas of individual stable isotope values of δ^13^C and δ^15^N for Charcotia obesa and its isotopic niche, grouped per depth. Different depths are indicated by different colours; the sample number per species is also provided.

### Tables

**Table 1S.** Sample size (n) and mean ± SD δ13C and δ15N values calculated for *Charcotia obesa*. Corrected values (corr) are adjusted for locations. Values are grouped according to sampling depth.

| Species | Depth | n | $\delta^{13}\text{C}$ (‰) | $\delta^{15}\text{N}$ (‰) | $\delta^{13}\text{C}_{\text{corr}}$ (‰) | $\delta^{15}\text{N}_{\text{corr}}$ (‰) |
| --- | --- | --- | --- | --- | --- | --- |
| *C. obesa* | <50 m | 115 | -23.2±1.0 | 11.2±1.1 | -23.6±0.5 | 10.7±0.6 |
| | 50-100 m | 29 | -24.4±0.5 | 9.7±0.8 | -24.0±0.3 | 10.5±0.5 |
| | 100-150 m | 34 | -24.1±0.2 | 10.3±0.5 | -23.8±0.3 | 10.9±0.6 |
| | >150 m | 25 | -24.9±0.3 | 9.8±0.5 | -23.9±0.3 | 10.6±0.5 |

**Table 2S.** Results of pair-wise Dunn tests with Bonferroni corrections assessing the influence of depths on corrected stable isotope values of carbon and nitrogen for *Charcotia obesa*. Significant p values (<0.05) are indicated in bold.

| Species | Depths | p-value | p-value |
| --- | --- | --- | --- |
| | | $\delta^{13}\text{C}_{\text{corr}}$ (‰) | $\delta^{15}\text{N}_{\text{corr}}$ (‰) |
| *C. obesa* | <50 m – 50 to 100 m | **<0.001** | **0.041** |
|  | <50 m – 100 to 150 m | 0.112 | 0.808 |
|  | <50 m – >150 m | 1 | 1 |
|  | 50 to 100 m – 100 to 150 m | 0.599 | **0.004** |
|  | 50 to 100 m – >150 m | 0.065 | 0.443 |
|  | 100 to 150 m – >150 m | 1.000 | 0.983 |

## Notes

### Competing Interest Statement

The authors have declared no competing interest.

